# Reprogramming Human Inflammatory Macrophages in Symptomatic Carotid Stenosis: Potential Mechanisms for Stabilisation of Atherosclerotic Carotid Plaques

**DOI:** 10.1101/2024.06.11.598440

**Authors:** Klaudia Kocsy, Sumeet Deshmukh, Shah Nawaz, Ali N Ali, Sheharyar Baig, Joyce S Balami, Arshad Majid, Endre Kiss-Toth, Sheila Francis, Jessica Redgrave

## Abstract

**Background:** Inflammation is a precursor to atherosclerotic plaque destabilisation, leading to ischaemic events such as stroke. Macrophage phenotypes can be altered by the microenvironment, and certain anti-inflammatory agents may, therefore, stabilise plaques and reduce the risk of recurrent ischaemic events.

**Methods:** Thirteen carotid plaques were obtained from stroke/ Transient Ischaemic Attack (TIA) patients undergoing carotid endarterectomy. An immunofluorescence stain was used to identify common macrophage markers (pan macrophage: CD68, pro-inflammatory: CD86, anti-inflammatory: MRC1), and a novel analysis technique was used to measure the prevalence of macrophage phenotypes in carotid plaques in relation to other histological features of instability.

An *in vitro* model of human blood-derived macrophages was also developed to evaluate the effect of statins and glucocorticoids on macrophage-specific markers using RT-qPCR, Western Blot and immunofluorescence stain. The physiological effect of dexamethasone was further evaluated on macrophages and human carotid plaques cultured *ex vivo*.

**Results:** The macrophage population (CD68+) in the carotid plaques was dominated by “double-positive” (CD86+MRC1+) macrophages (67.8%), followed by “M1-like” (CD86+MRC1-) (16.5%), “M2-like” (CD86-MRC1+) (8.7%) and “double-negative” (CD86-MRC1-) (7.0%) macrophages. M1-like macrophages were more prevalent in unstable plaque sections than stable ones (p=0.0022).

Exposure to dexamethasone increased macrophage *MRC1* gene expression *in vitro* and *ex vivo*. Dexamethasone also reduced Oxidised Low-Density Lipoprotein Receptor 1 (*OLR1*) gene and protein expression, leading to a decreased ox-LDL uptake in foam cell assays. This was, in turn, associated with reduced lipid uptake in macrophages, as shown by Oil Red O staining.

**Conclusions:** Human macrophages may be “switched” to a less inflammatory phenotype by exposure to clinically relevant concentrations of glucocorticoid, potentially mediated by a reduction in Oxidised LDL uptake. This effect was not observed following macrophage exposure to statins. Glucocorticoids may have a future role in preventing ischaemic events in patients with advanced atherosclerosis.

**Graphical Abstract:** 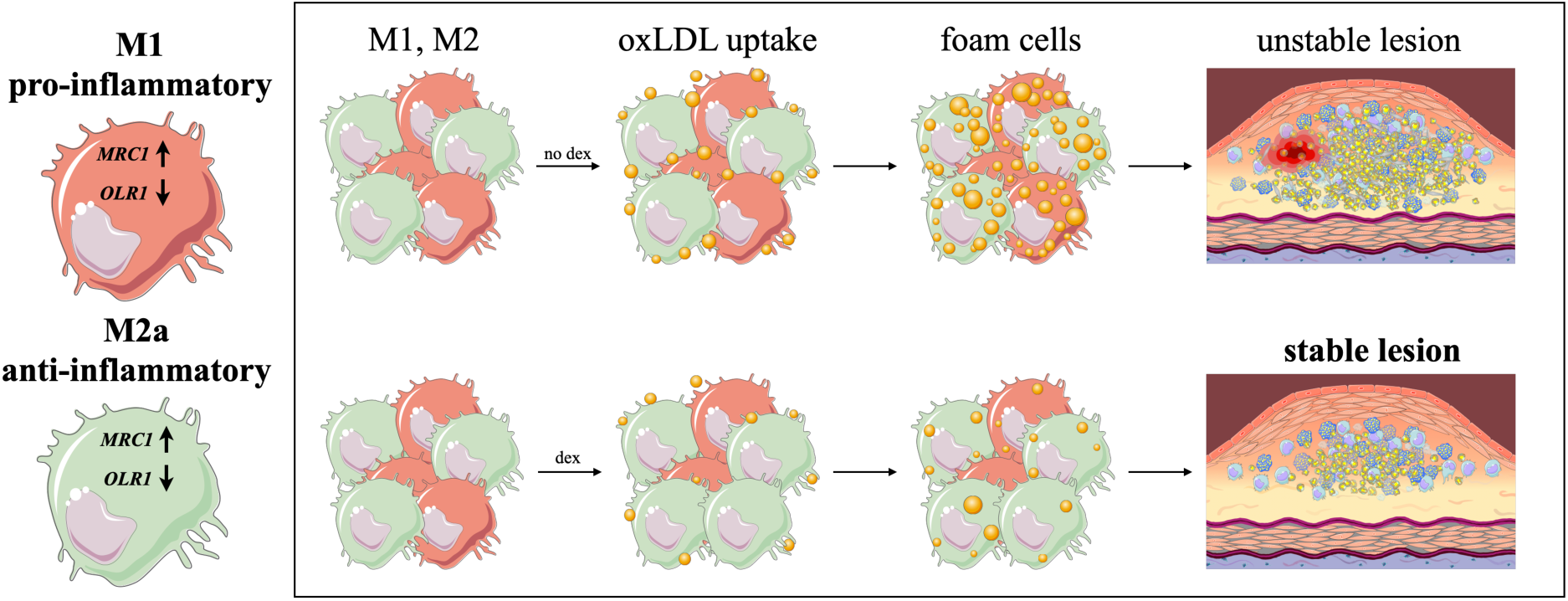

**Highlights:**

- A high prevalence (68% in this study) of carotid plaque macrophages express both pro-inflammatory (CD86) and anti-inflammatory (MRC1) markers. These may represent a novel macrophage population.
- Human macrophages may be “reprogrammed” to a less inflammatory phenotype following exposure to glucocorticoids.
- Dexamethasone increased *MRC1* and decreased *OLR1* expression in macrophages derived from human blood samples *in vitro* and in cells derived from cultured human carotid plaque tissue *ex vivo.* This was associated with reduced oxLDL uptake and reduced lipid accumulation in the macrophages.
- Dexamethasone has the potential to stabilise carotid atherosclerotic plaques in humans.

## Introduction

Atherosclerosis is a systemic inflammatory disease during which lipoproteins, cholesterol, calcium and apoptotic cells build up to form a lipid-rich plaque within an artery [1]. Large artery atherosclerosis is the most frequent cause of ischaemic stroke, and patients with a high atherosclerotic plaque burden are at increased risk of early recurrent ischaemic events [2]. Macrophages, lipoproteins, and foam cell formation are critical components of atherosclerotic plaque formation, progression and instability [3, 4]; therefore, targeting these could provide a potential novel treatment modality for stroke patients.

Macrophages are essential in regulating lipid metabolism as they take up low-density lipoproteins (LDL), very-low-density lipoprotein (VLDL) and oxidized low-density lipoprotein (oxLDL) via scavenger receptor-mediated pathways, phagocytosis or micropinocytosis [5]. Macrophages can also help to eliminate cholesterol. If excess cholesterol is not removed, it can lead to foam cell formation and plaque progression [5]. A large symptomatic carotid plaque study of 526 plaques found a high prevalence of macrophage infiltration, lipid core and foam cells [6]. It led to the derivation and validation of a reproducible histology technique to measure overall plaque instability [7, 8].

Tissue macrophage cell populations are heterogeneous, and there is increasing evidence that macrophages can “switch” phenotypes between proinflammatory and anti-inflammatory subsets in the developing atherosclerotic plaque [4]. Broadly, the classically activated, proinflammatory macrophages (usually referred to as M1) promote atherogenesis and enhance plaque instability as they secrete proinflammatory cytokines such as IL-1β, IL-7, IL-8, IL-12 and IL-6. In contrast, alternatively-activated, anti-inflammatory macrophages (usually referred to as M2) have regulatory and wound-healing properties, including collagen synthesis and promoting tissue repair [4]. They produce anti-inflammatory cytokines and have a potential role in the maintenance of plaque stability due to their immunoregulatory functions [9].

In atherosclerotic plaques, macrophages have a specific spatial distribution, significantly impacting plaque stability [10]. For example, proinflammatory macrophages are often found in the shoulder region of the plaque, where cap disruption most often occurs, while anti-inflammatory macrophages are primarily found in the adventitia layer [10]. In another study of 65 carotid plaques, proinflammatory markers (e.g. IL6, MMP9) were dominant in symptomatic plaques, while in asymptomatic plaques, anti-inflammatory macrophage markers (CD163, MRC1) were dominant [11].

A single macrophage can express both proinflammatory (e.g. CD86) and anti-inflammatory (e.g. MRC1) markers [12]. These may represent macrophages transitioning from one phenotype to another, perhaps in response to their microenvironment. However, their prevalence and potential association with plaque stability have not been studied in detail before.

Macrophages and other inflammatory cells can “polarise” in response to their microenvironment and initiate certain inflammatory signalling pathways, e.g. cytokine signalling, glucocorticoid receptor (GR) signalling or lipopolysaccharide (LPS) signalling, which can then trigger biological responses [13]. Therefore, it is plausible that exposure to certain pharmacological agents, especially those purported to have anti-inflammatory properties, like 3-hydroxy-3-methylglutaryl coenzyme A (HMG-CoA) reductase inhibitors (statins) or glucocorticoids, may affect their functional phenotype [4, 14].

Statins are widely used in primary and secondary cardiovascular risk modification in humans as they reduce the risk of cardiovascular events by decreasing LDL levels in the bloodstream [15]. Statins are also purported to have anti-inflammatory effects as they increase macrophage apoptosis through the c-Jun N-terminal kinase (JNK) pathway and reduce macrophage colony-stimulating factor-mediated proinflammatory activities [15]. This might be one mechanism for the observation that statins can lead to the regression of intraplaque lipids in longitudinal plaque imaging studies [15].

Glucocorticoids are another class of drugs which are exploited for their anti-inflammatory effects. These essential stress hormones play a role in inflammation, homeostasis, cell proliferation and development [16]. Glucocorticoids are used widely to treat immune system diseases and their effect is highly diverse; they can increase apoptosis of inflammatory cells, decrease cytokine production, activate histone deacetylase and increase gene transcription [13]. Dexamethasone, in particular, has been shown to inhibit macrophage proliferation and suppress foam cell formation *in vitro* [17]. The concentration of dexamethasone used in both *in vitro* and *in vivo* studies has been highly variable [17], [18], [19], [20]. For the current study, clinically relevant concentrations of anti-inflammatory agents (i.e. those reached in the blood when the agents are administered therapeutically to humans) were determined from dose-dependent pharmacokinetic studies [21] [22] (Supplementary) and deployed.

This study aimed to explore whether disease-enhancing macrophage phenotypes can be modulated pharmacologically, thereby providing a mechanism by which future therapies could reduce the risk of plaque destabilisation and recurrent ischaemic events.

Our objectives were:

1. To determine the prevalence of macrophage subsets in unstable and stable carotid plaque regions.
2. To determine the effect of statins and glucocorticoids on macrophages derived from blood from healthy human volunteers and patients with recent stroke using an *in vitro* model.
3. To further assess the effect of dexamethasone on macrophage phenotype and oxidised LDL uptake in human carotid plaques cultured *ex vivo*.

To our knowledge, this is also the first study addressing the direct role of glucocorticoids on human carotid plaque tissue, which has been excised from patients and cultured *ex vivo*.

Our data revealed that dexamethasone “reprogrammed” macrophages to a less inflammatory state, this was accompanied by reduced oxidised LDL uptake and reduced lipid accumulation in the macrophages.

## Methods

A detailed description of the methods is provided in the Supplementary Methods.

All experiments were performed in accordance with United Kingdom legislation under the Human Tissue Act 2004. Carotid tissue and blood samples were collected from The Northern General Teaching Hospitals in Sheffield (REC reference 14/SC/0147).

### Histological Features in Atherosclerotic Plaques

Patients (n=13) with a recent stroke or TIA and greater than 50% stenosis of the ipsilateral internal carotid artery undergoing carotid endarterectomy (CEA) were recruited for this study and atherosclerotic plaque and/or peripheral blood were collected.

Eleven good-quality human carotid plaques (2 were excluded because of excessive fragmentation) were divided into up to 5 sections (A-E) depending on size, resulting in 46 sections from 11 plaques [8] (Supplementary Table 1). The sections were taken from the bifurcation and 3 mm on either side to capture the majority of unstable plaque features where they were to be found anywhere along the length of the plaque as per the sectioning technique of Lovett et al [8]. Plaques were prepared and stained with H&E, Verhoeff-van Gieson stain, CD3 and CD68. Histological features (lipid core, fibrous cap, intraplaque haemorrhage, macrophages, lymphocytes, calcification) were measured using previously published detailed semiquantitative scales, leading to a composite assessment of “overall instability” [8]. Plaques were also graded according to the American Heart Association classification for the degree of advancement of atherosclerosis [23, 24].

### Macrophage Subsets in atherosclerotic plaques

Macrophage phenotypes from the shoulder regions (from 23 sections from 11 plaques) were determined with immunofluorescence (IF) microscopy. Briefly, a single IF stain with a pan macrophage polarisation marker (CD68) was used alongside a dual IF stain with M1 (CD86) and M2 markers (MRC1). The regions of positive staining for CD68 were “overlain” with the dual stained section, and the prevalence of CD86+ and MRC1+ macrophages were quantified by Fiji 2.0 [25].

### Reprogramming Human Macrophages

An *in vitro* model was developed based on a previously established method [26] to characterise macrophages and their response to different pharmaceutical agents (**Figure 1**). Briefly, human monocyte-derived macrophages (MDMs) were differentiated from peripheral blood monocytes from healthy volunteers and stroke patients’ blood samples. MDMs were then polarised towards either M^LPS+INFγ^ or the M^IL4^ phenotype. The polarised hMDMs were treated with atorvastatin, simvastatin, dexamethasone, prednisolone or hydrocortisone in concentrations known to equate to plasma concentrations in human subjects [21] [22].

**Figure 1:**
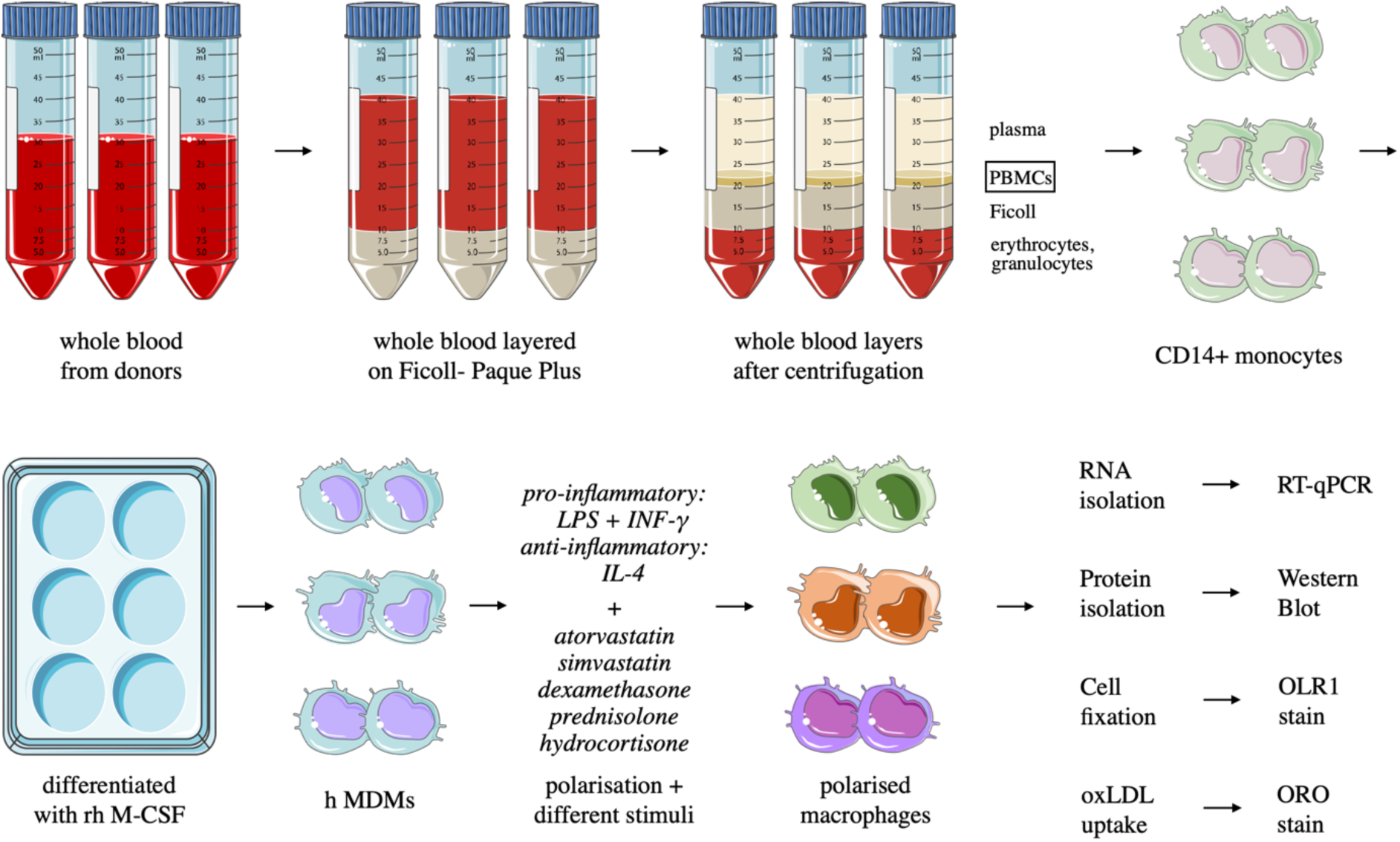
Workflow of in vitro cell work. *PBMCs were isolated from whole blood, and the CD14+ monocytes were selected and then differentiated into monocyte-derived macrophages with rh M-CSF. Polarised human MDMs were treated with atorvastatin, simvastatin, dexamethasone, prednisolone or hydrocortisone in different clinically relevant concentrations. Gene expression analysis was carried out in the differentiated hMDMs to test for macrophage-specific genes and scavenger receptors. Western Blot analysis and immunofluorescence stain (IF) were performed on dexamethasone-treated MDMs to validate changes in the expression of OLR1 protein. The physiological effect of dexamethasone was also tested on oxLDL uptake by macrophages*.

For the *in vitro* studies, RNA or protein was isolated from drug-treated polarised hMDMs differentiated from either healthy volunteers (n=9) or stroke patients’ (n=7) peripheral blood mononuclear cells (PBMCs). Gene expression analysis with RT-qPCR was conducted to test for macrophage-specific genes and scavenger receptors (*MRC1*, *OLR1*, *SCARB1*, *MSR1*, *CD36*). Western Blots (7 healthy volunteers) and IF stains (5 healthy volunteers) were performed to verify the change of OLR1 expression at a protein level. A functional test for lipid accumulation was conducted to assess the oxLDL accumulation in dexamethasone-treated hMDMs using Oil Red O staining (on blood from 5 healthy volunteers and 7 stroke patients).

The effect of dexamethasone was further tested *ex vivo* on human carotid plaque tissue excised from 4 stroke/TIA patients. After carotid endarterectomy, plaque tissue was divided into 2 main parts. The first part was dissected into smaller pieces and cultured with and without dexamethasone for 3 days. Then, a single-cell suspension was made and gene expression analysis was performed. The second part of each plaque was divided into groups (day 0, day 3 untreated and day 3 dexamethasone-treated) and IF stained. The CD68 pan macrophage marker was used along with the MRC1 (M2) marker, or the CD68 marker was paired with the OLR1 antibody. The fluorescence intensity of MRC1 and OLR1 protein of the CD68+ macrophages was quantified by FIJI Image J [25].

### Statistics

All statistical analyses were performed with the Prism 9 program (GraphPad Software, Inc.) at a significance level of 95% (*P*<0.05). All statistical analyses are detailed within the corresponding figure legends.

## Results

To characterise plaque macrophage phenotypes, carotid plaques (n=13) were collected immediately following CEA from patients with confirmed stroke and with a greater than 50% stenosis of the internal carotid artery. Four additional plaques were collected for the ex vivo work.

Of the 17 patients (69% male) who donated their carotid plaques, all had had a TIA or an ischaemic stroke (on the ipsilateral side) within the previous 2 months (mean age: 72.64, SD: 9.98). Further patient demographic and clinical data are summarised in Supplementary Table 1.

The sufficiently intact plaque specimens (n=11) were divided into regions (n=46 analysed sections) and classified histologically (Supplementary Table 2). From the areas examined, 26.1% were designated as “definitely unstable” on histology, 26.1% as “probably unstable”, 26.1% as “probably stable”, and 21.7% as “definitely stable” according to Lovett et al. grading scale [8] **(Figure 2:A**)

**Figure 2:**
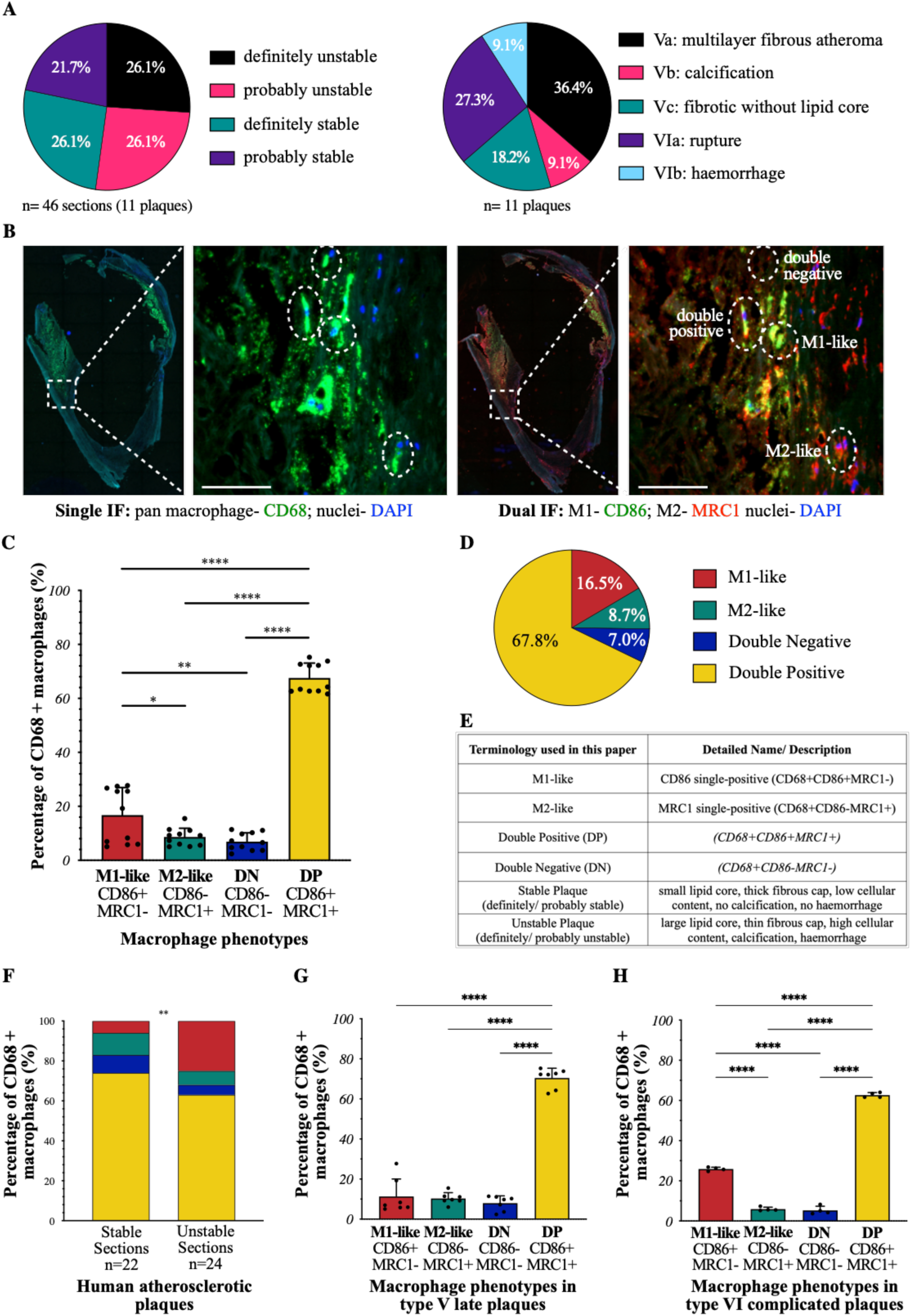
Histological features and different macrophage phenotypes of human carotid plaques. *Human carotid plaque classification is based on overall stability (**A/1**) [7, 8] and according to the American Heart Association grading scale (**A/2**) [23, 24] based on 2 independent observers (n=11 patients, 46 sections). **B:** IF stained FFPE human carotid atherosclerotic plaques from stroke patients. Scale bar= 50 µm **B/1, B/2:** stained for pan macrophage marker: CD68 (green); **B/3, B/4:** simultaneously stained for M1: CD86 (green) and for M2: MRC1 (red). **B/1, B/3:** IF stained serially next, entire carotid sections. **B/2, B/4:** enlarged regions of the previously identified shoulder regions of the single and dual IF stained sections. Dotted circles indicate the 4 main macrophage types: double-positive, double-negative, M1-like and M2-like macrophages. **C, D**: The regions of positive staining for CD68 were overlaid with the dual stained section, and the prevalence of CD86+ and MRC1+ macrophages were quantified by Image J and presented on **C:** bar graph and **D:** pie chart. The double-positive macrophage ratio is significantly higher than the other CD68+ cell groups. The proportion of M2-like and double-negative macrophages is similar, while M1-like macrophages account for 16.5% of all CD68+ cells.* **E:** *Summary of the terminology used throughout this paper.* **F:** *Macrophage phenotype distribution in stable and unstable carotid plaques. Data is plotted from 24 unstable and 22 stable carotid sections and analysed with a Chi-square test to determine the difference between groups.* **G, H:** *Percentage of different macrophage phenotype populations in AHA-classified plaque types*, *determined with single (CD68) and dual (CD86, MRC1) IF stains.* **G:** *In* type V-late plaques (n=7) *the ratio of CD86 single-positive (CD68+CD86+MRC1-), MRC1 single-positive (CD68+CD86-MRC1+) and double-negative (CD68+CD86-MRC1-) macrophages is similar, around 10%, while the percentage of double-positive (CD68+CD86+MRC1+) macrophages is significantly higher (7*0*%).* **H:** *In* type VI-complicated plaques (n=4), *the CD86 single-positive (CD68+CD86+MRC1-) macrophage ratio (approximately 2*6*%) is higher than the MRC1 single-positive (CD68+CD86-MRC1+) (*6*%) or double-negative (CD68+CD86-MRC1-) (*5*%) macrophages. The double-positive (CD68+CD86+MRC1+) macrophage proportion in* these *plaque types is around 6*3*%, significantly higher than the other macrophage groups.* **All:** *Data follow normal (Gaussian) distribution, analysed with ordinary one-way ANOVA with Tukey’s multiple comparison test and the graphs presented with mean + SD*. *n=11 individual patient samples; p-value: *‹0.0332, ** ‹0.0021, **** ‹0.0000*.

### Macrophage phenotype characterisation in carotid plaques

The plaque shoulder regions (defined as the region where the fibrous cap meets the normal adventitia wall) were identified on H&E-stained sections from the good-quality plaques. These areas were designated “regions of interest” (23 ROI from 11 carotid plaques), as the shoulder region is regarded as the most vulnerable area of the plaque, where the cap is often at its thinnest [27].

Our novel, dual IF staining **(Figure 2: A)** identified 4 different macrophage phenotypes: “M1-like” (CD86 single-positive: CD68+CD86+MRC1-), “M2-like” (MRC1 single-positive: CD68+CD86-MRC1+), “double-positive” (CD68+CD86+MRC1+) and “double-negative” (CD68+CD86-MRC1-) macrophages **(Figure 2: B)**.

Of the 23 analysed plaque sections, macrophages were abundant in 22 (n=22 sections from n=11 human plaques). Double-positive macrophages were the most prevalent overall (67.8%) **(Figure 2:C, D)**. M1-like macrophages accounted for 16.5%, while M2-like macrophages comprised only 8.7%. Double-negative macrophages represented 7.0% of all macrophages.

The proportion of different macrophage subsets in stable and unstable carotid plaque sections (as defined by Lovett et al [7, 8]) was evaluated **(Figure 2: F)**. The proportion of M1-like macrophages was greater in the unstable plaque sections (p= 0.0022), whereas M2-like, double-negative and double-positive macrophages were more prevalent in the stable plaque sections. Double-positive and double-negative macrophage phenotypes were found to a similar degree in both stable and unstable carotid plaque sections.

Results of the AHA grading of plaques [23, 24] are detailed in Supplementary Table 4. There were 7 AHA type V (late) plaques and 4 AHA type VI (complicated) plaques.

The percentage of double-positive (M1 and M2 hybrid) macrophages was the highest of all CD68+ macrophages (70% in type V plaques and 63% in type VI plaques). The percentage of M1-like macrophage population was 11% of all CD68+ macrophages in type V plaques, while it was significantly higher (26%) in the complicated type VI plaques (p= 0.0003). M2-like and double-negative macrophage populations were similarly represented in both AHA plaque types (M2-like in type V plaque: 10%; M2-like in type VI plaque: 6%; DN in type V plaque: 8%; DN in type VI plaque: 5%).

On review of the literature, these double-negative (CD68+CD86-MRC1-) and double-positive (CD68+CD86+MRC1+) macrophage subtypes were also identifiable in the cite-seq (cellular indexing of transcriptomes and epitopes: simultaneous sequencing of cell surface protein and transcriptomic data) dataset published by Fernandez et al. [28] **(Figure 3)**. In their dataset, 254 plaque macrophages were sequenced, and after re-analysing their published dataset, we have identified that 143 cells (56%) were M1-like, 9 cells (4%) were M2-like, 34 (13%) were double-negative while 68 (27%) cells were double-positive macrophage subtypes **(Figure 3)**. We have further analysed these subgroups in a cluster analysis (**Figure 3: B)**. The double-positive macrophages are clustered together, implying that they are a distinct phenotype, while the double-negative macrophages are scattered across all macrophage groups.

**Figure 3:**
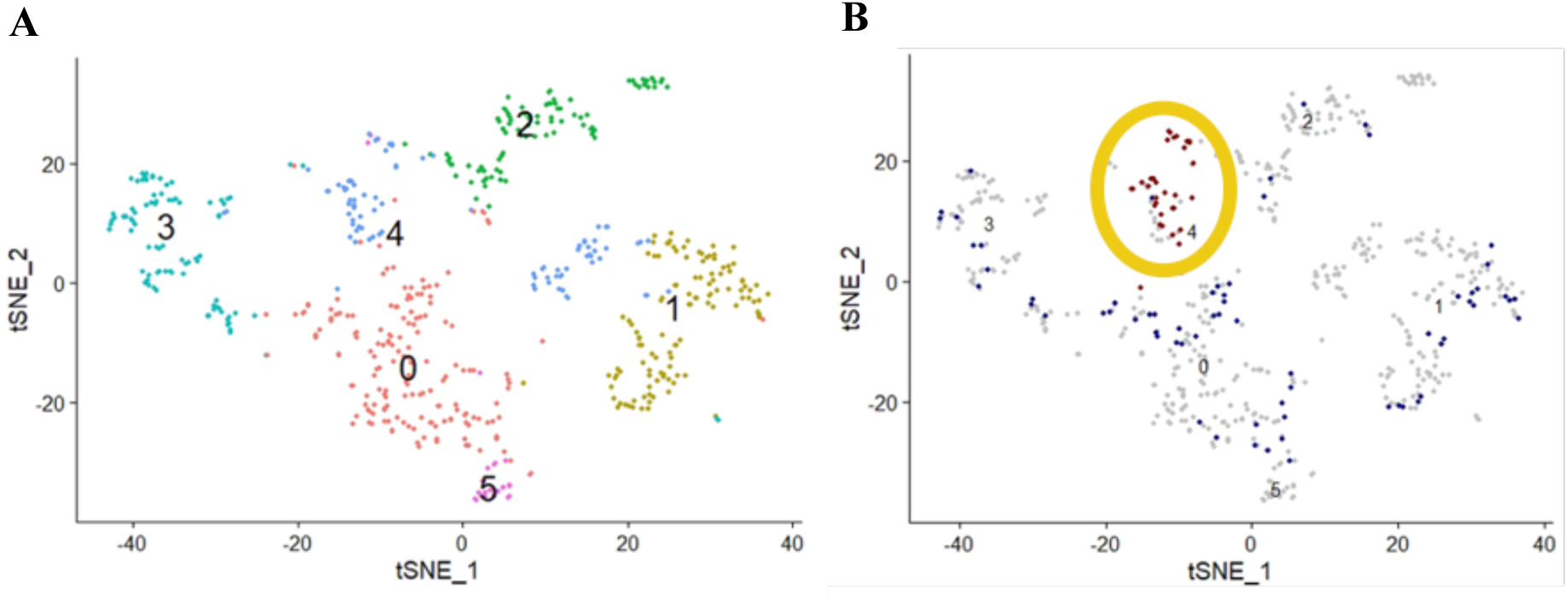
Identified double-negative and double-positive macrophages in the cite-seq dataset from Fernandez et al. [28]. *Cluster assignment of 254 plaque macrophages visualised with t-SNE plot (t-Distributed Stochastic Neighbor Embedding). **A**: Cluster assignments of all plaque macrophages are defined by Louvain clustering and denoted as distinct colours. **B**: Double positive and double negative plaque macrophages are highlighted. 27% of the plaque macrophages are identified as DP (yellow circled area), while 13% are deemed DN (blue dots). (DN= double-negative, DP= double-positive)*.

### Dexamethasone promotes polarisation towards an anti-inflammatory state

Since anti-inflammatory glucocorticoids may have the capacity to modulate the development of atherosclerosis [13], we investigated whether glucocorticoids upregulate the Mannose Receptor C-type 1 gene (*MRC1*), an anti-inflammatory marker in polarised macrophages [29].

The effect of 3 different glucocorticoids on macrophages was tested using equivalent and clinically relevant concentrations of dexamethasone (98 ng/ml), prednisolone (5.2 µg/ml) and hydrocortisone (2.8 µg/ml) (glucocorticoid dose equivalency based on information from the British National Formulary). There was a similar (2-fold) increase in the *MRC1* gene expression seen following exposure to both dexamethasone (M^LPS+INFγ^: p= 0.0043, M^IL4^: p= 0.0254) and prednisolone (M^LPS+INFγ^: p= 0.0014, M^IL4^: p= 0.0155) (**Figure 4: A/1, A/2**).

**Figure 4:**
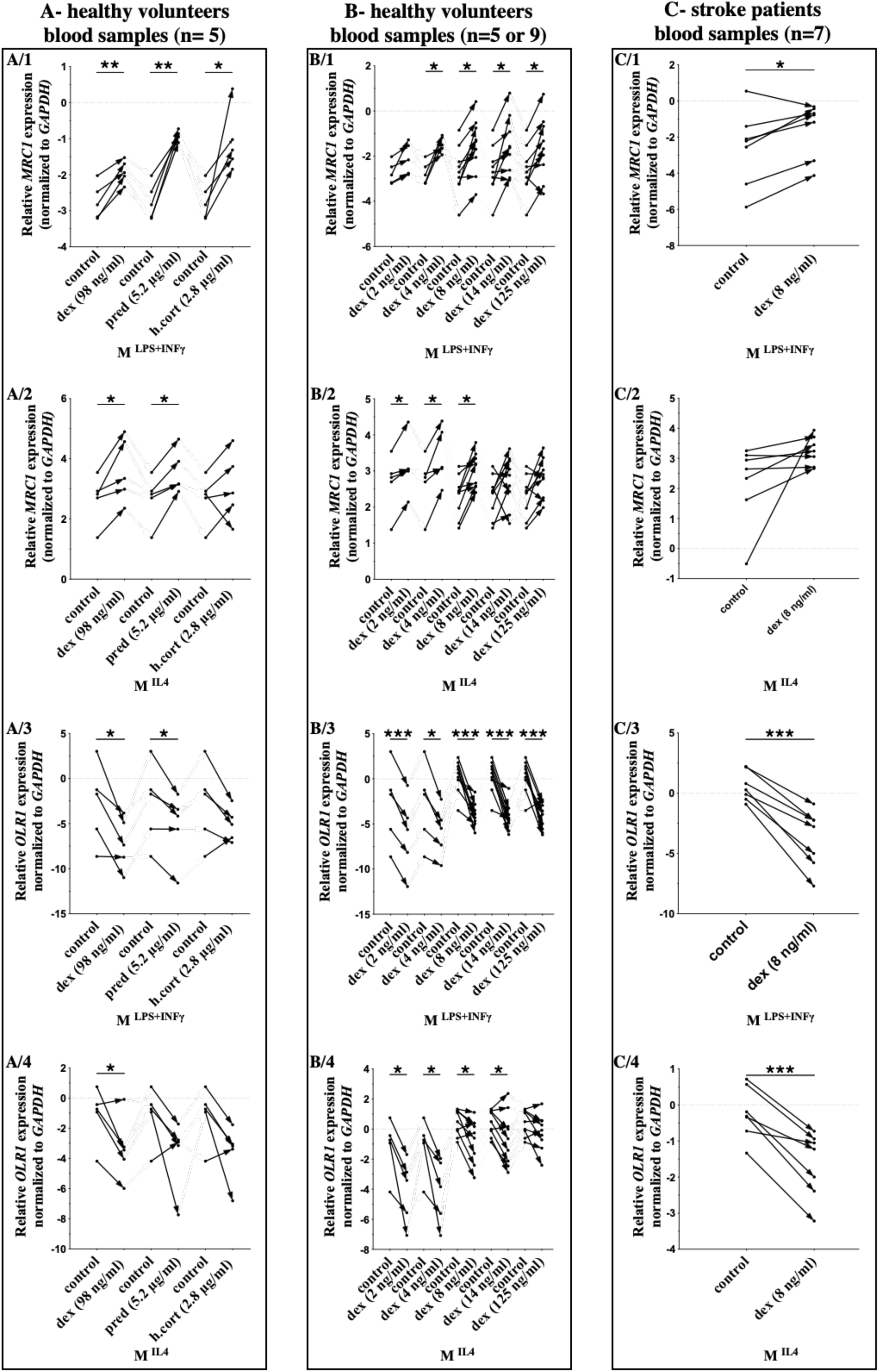
Different glucocorticoids promote polarisation towards an anti-inflammatory state and decrease OLR1 gene expression in human macrophages. *Human M^LPS+INFγ^ (1^st^, 3^rd^ rows) and M^IL4^ (2^nd^, 4^th^ rows) macrophages were treated with either dexamethasone, prednisolone or hydrocortisone, and RT-qPCR was performed. Generally, MRC1 (1^st^, 2^nd^ row) gene expression was increased, and OLR1 (3^rd^, 4^th^ row) expression was decreased following all glucocorticoid treatment. **A:** MDMs, isolated from the blood of 5 healthy volunteers (n=5), were treated with dexamethasone (98 ng/ml), prednisolone (5.2 µg/ml) or hydrocortisone (2.8 µg/ml). **B:** Macrophages isolated from blood samples of healthy volunteers (n=5 or 9) were treated with dexamethasone in lower concentrations (2 and 4 ng/ml) as well as clinically relevant concentrations (8, 14, 125 ng/ml). **C:** Macrophages from stroke patients (n=7) blood samples were treated with 8 ng/ml dexamethasone, and downregulation of OLR1 gene expression was highly significant following dexamethasone treatment **(C/3, C/4)**. **All:** Data is individually listed and normalised to the GAPDH housekeeping gene. Data follow normal (Gaussian) distribution and are analysed with a two-tailed paired t-test. The direction of change in expression is shown with an arrow for each sample tested. The dotted line indicates a 0-expression level. p-value: * ‹0.0332, ** ‹0.0021, *** ‹0.0002*.

The dose-dependent effect of dexamethasone was further tested on human polarised M^LPS+INFγ^ and M^IL4^ macrophages isolated from healthy adult volunteer blood samples (**Figure 4: B**). *MRC1* gene expression was increased after exposure to dexamethasone at lower doses than are normally used in clinical practice (2, 4 ng/ml) as well as at all clinically relevant concentrations of dexamethasone (8, 14, 125 ng/ml) (**Figure 4: B/1, B/2**). There was a significant change following exposure to 4 and 8ng/ml dexamethasone treatment concentrations in both M^LPS+INFγ^ and M^IL4^ macrophage subtypes.

Finally, macrophages derived from the blood of stroke survivors also showed an increase in *MRC1* expression (M2 marker) following exposure to dexamethasone (**Figure 4: C/1, C/2**). In summary, these data show that macrophages can become polarised towards a less inflammatory phenotype following exposure to glucocorticoids (dexamethasone and prednisolone).

### Dexamethasone decreases OLR1 expression

*OLR1* encodes for a low-density lipoprotein receptor that binds, internalises, and degrades oxLDL and has been associated with the progression of atherosclerosis [30]. Therefore, we investigated whether glucocorticoids downregulate the expression of OLR1 as assessed by RT-qPCR, WB and IF stain. *OLR1* gene expression was decreased following 24-hour treatment with all three glucocorticoids (**Figure 4: 3^rd^ and 4^th^ row**), but dexamethasone significantly decreased *OLR1* expression in both M^LPS+INFγ^ and M^IL4^ polarised cells (**Figure 4: A/3, A/4**).

A dose-dependent effect of dexamethasone was sought separately on M^LPS+INFγ^ and M^IL4^ macrophages isolated from the blood of healthy volunteers and stroke patients. *OLR1* gene expression decreased after dexamethasone exposure at all the tested concentrations (2, 4, 8, 14, 125 ng/ml) (**Figure 4: B/3, B/4**).

Dexamethasone significantly decreased *OLR1* gene expression in macrophages isolated from blood from stroke patients (M^LPS+INFγ^: p= 0.0003, M^IL4^: p= 0.0009) (**Figure 4: C/3, C/4**).

This decrease in OLR1 was verified using Western Blot analysis (**Figure 5: A, B, G)** and immunofluorescence staining (**Figure 5: C, D, H)**. OLR1 protein expression, normalised to GAPDH, was reduced following 24-hour exposure to dexamethasone in both M^LPS+INFγ^ (8 ng/ml: p= 0.0003, 14 ng/ml: p= 0.0017) (**Figure 5: A)** and M^IL4^ (8 ng/ml: p= 0.0232, 14 ng/ml: p= 0.0813) (**Figure 5: B)** macrophage subtypes.

**Figure 5:**
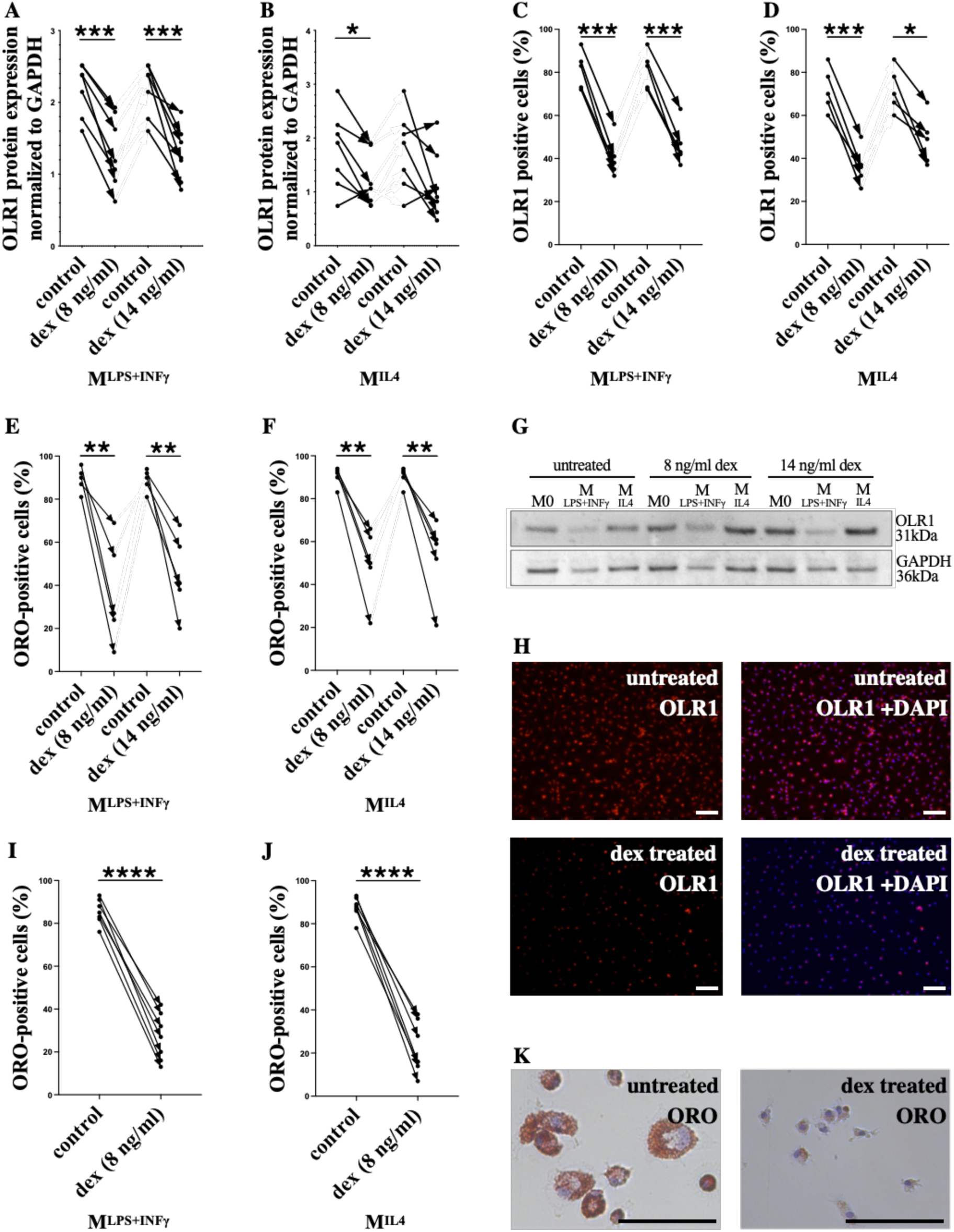
OLR1 protein expression and oxLDL binding capability of human MDMs were reduced following exposure to dexamethasone. *Human* M^LPS+INFγ^ *(A, C, E, I) and M^IL4^ (B, D, F, J) polarised macrophages were treated with dexamethasone (8, 14 ng/ml). **A, B, G:*** OLR1 *protein expression was significantly decreased, determined by Western Blot. The expression of* OLR1 *protein was normalised to* GAPDH *housekeeping protein and plotted from healthy volunteers (n=5). **G**: A representative Western Blot is shown. **C, D, G**:* OLR1 *protein expression was significantly decreased by dexamethasone treatment, measured by IF (n=7).* OLR1 *positive cells were counted and compared to the maximum cell number based on the DAPI IF signal and quantifications performed on an average of 5 fields of view. **H**: Representative immunofluorescence images of non-polarised hMDMs with OLR1 (red) and nuclei (DAPI, blue) markers. Dexamethasone-treated hMDMs are shown at the bottom*. ***E, F, I, J, K:*** *Lipid accumulation was tested by adding 25µg/ml oxLDL and assessed by Oil Red O stain. The percentage of positive cells was counted, and quantifications were performed on an average of 5 fields of view. Data is plotted from 5 healthy volunteers (E, F; n=5) and 7 stroke patients (I, J; n=7). Lipid binding was reduced in hMDMs treated with dex, which is highly significant in cells isolated from stroke patients (I, J). **K**: Representative images of untreated (left) and dexamethasone-treated (right) hMDMS after ORO stain. **H, K**: Scale bar 100 µm. **All**: Data follow normal (Gaussian) distribution and are analysed with a two-tailed paired t-test. The direction of change in expression is shown with the treatment as an arrow for each sample tested. P-value: * ‹0.0332, ** ‹0.0021, *** ‹0.0002 **** ‹0.0001*.

The change in OLR1 protein expression after 24h dexamethasone treatment was further validated using IF staining for surface receptors. The percentage of OLR1-positive cells reduced by an average of 35% after 24 hours of exposure to dexamethasone (**Figure 5: C, D)**.

These findings suggest that dexamethasone-treated macrophages might bind less oxLDL, which could, in turn, reduce the formation of atherosclerosis in the vessel wall.

The effect of dexamethasone on *SCARB1*, *MSR1* and *CD36* genes was also tested with RT-qPCR, and results are shown in Supplementary Figure 3.

### The ox-LDL binding capability of polarised macrophages is decreased by dexamethasone treatment

Following the finding of reduced OLR1 expression following dexamethasone exposure, the ox-LDL accumulation in dexamethasone-treated hMDMs was assessed using Oil Red O (ORO) staining.

In both M^LPS+INFγ^ **(Figure 5: E, I)** and M^IL4^ **(Figure 5: F, J)** cells, after 8 and 14 ng/ml dexamethasone treatment, the percentage of ORO-positive cells was significantly reduced (in healthy volunteer blood-derived macrophages by an average of 44% **(Figure 5: E, F)**, and in stroke patients blood-derived macrophages by an average of 62% **(Figure 5: I, J)**.

These findings suggest that dexamethasone can reduce the oxLDL binding capacity of macrophages, leading to reduced lipid accumulation and potentially less foam cell formation.

### The effect of dexamethasone on human carotid plaques, ex vivo

Finally, the effect of dexamethasone on MRC1 and OLR1 expression was investigated *ex vivo* on 4 freshly obtained human carotid plaques (i.e. tissue including all plaque cells, not just macrophages).

After carotid endarterectomy, plaque tissue was cultured with and without dexamethasone for 3 days (8 ng/ml), and the MRC1 and OLR1 mRNA and protein expressions were evaluated **(Figure 6)**.

**Figure 6:**
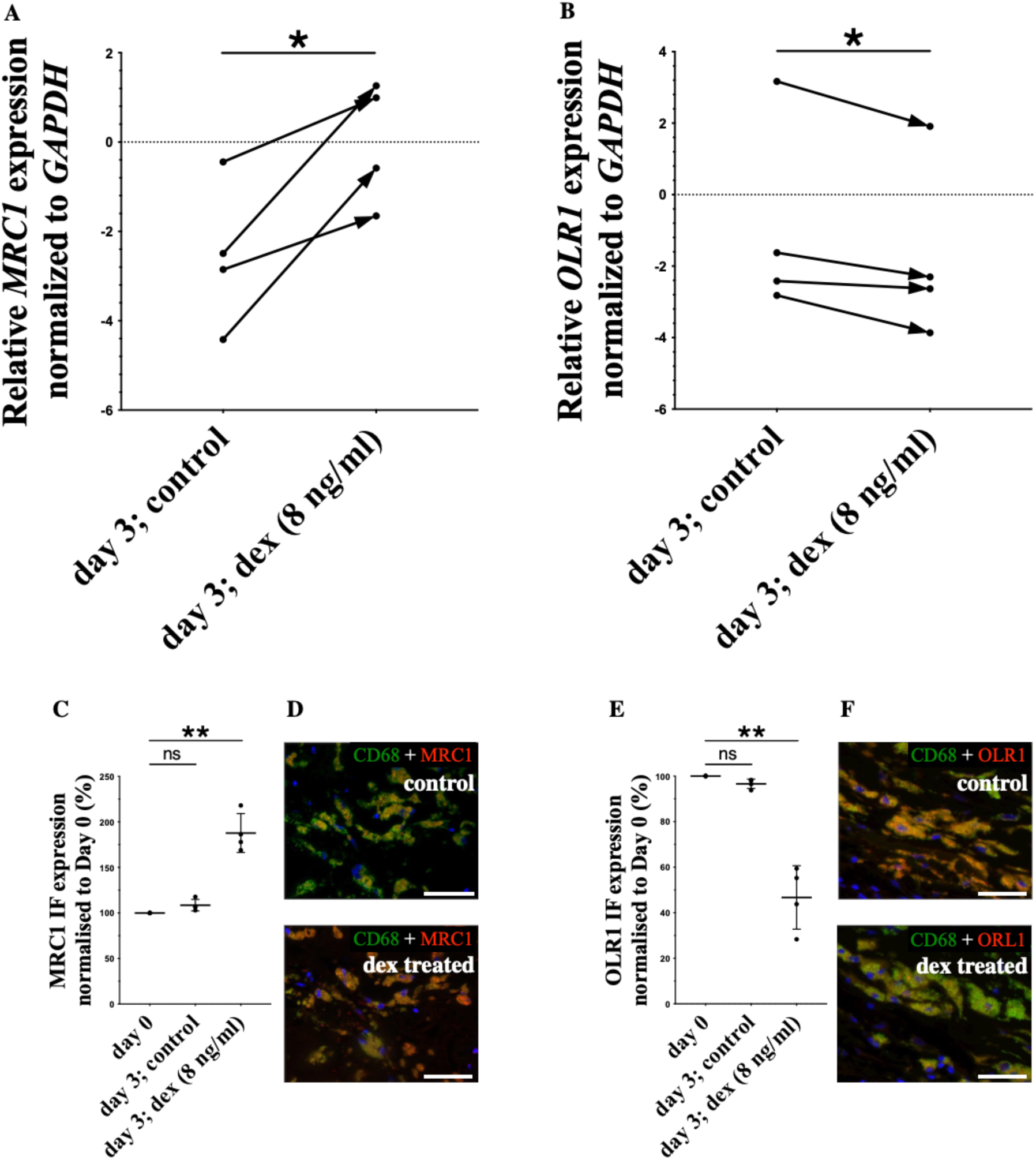
The effect of dexamethasone on ex vivo cultured human carotid plaques. *Carotid plaque tissue, removed by endarterectomy, was dissected into smaller pieces and cultured with or without dexamethasone. **A, B:** Single-cell suspension was made, and RT-qPCR was performed. MRC1 expression was increased (A), while OLR1 gene expression (B) was decreased in cells from human plaques following 3-day exposure to dexamethasone. Data is plotted from 4 carotid plaques (n=4 stroke/TIA patients), individually listed and normalised to the GAPDH housekeeping gene. Data follow normal (Gaussian) distribution and are analysed with a two-tailed paired t-test. The direction of change in expression is shown with the treatment as an arrow for each plaque. The dotted line indicates a 0-expression level. p-value: * ‹0.0332 **C-F:** Atherosclerotic plaques were divided into 3 main parts (day 0, day 3 untreated, day 3 dexamethasone-treated) and, after histological processing, were IF stained with CD68 and MRC1 (C, D), or the CD68 with the OLR1 antibody (E, F). The* MRC1 *and* OLR1 *IF expression of the CD68+ macrophages were measured and normalised to day 0 control. The* MRC1 *protein was significantly increased, and the* OLR1 *was significantly decreased ex vivo, compared to the untreated day 3 and day 0 control samples. Data is plotted from 4 carotid plaques (n=4 stroke/TIA patients), individually listed and analysed with one-way ANOVA with Dunnett’s multiple comparison test. P-value: ** ‹0.0021. **D, F:** Representative immunofluorescence images of ex-vivo treated macrophages are shown with CD68 (green) MRC1 or OLR1 (red) and nuclei (DAPI, blue) markers. Scale bar: 100 µm*.

RT-qPCR analysis confirmed that the *MRC1* RNA expression increased (p= 0.0377) **(Figure 6: A**), while *OLR1* expression was decreased (p= 0.0393) **(Figure 6: B**) following dexamethasone exposure.

The changes in MRC1 and OLR1 protein expression of CD68+ macrophages were assessed with IF staining **(Figure 6: C-F**). MRC1 protein levels significantly increased (p= 0.0062) **(Figure 6: C, D**), whilst OLR1 significantly reduced (p= 0.0077) **(Figure 6: E, F**) compared to the untreated day 3 and day 0 control samples.

These results suggest that dexamethasone, at clinically relevant concentrations, can also alter intra-plaque macrophages to become less inflammatory, thereby potentially offering a therapeutic role in plaque stabilisation.

### Statins do not have a significant effect on gene expression of hMDMs in vitro

Statins are widely prescribed to patients following ischaemic stroke and TIA as part of usual care in secondary prevention to reduce the likelihood of future ischaemic events. Statins have been shown to reduce cholesterol biosynthesis [31] as well as to modify LDL uptake by scavenger receptors on macrophages. Gene expression of macrophage-specific scavenger receptors after exposure to statins was therefore tested [32].

Polarised human blood monocyte-derived macrophages (M^LPS+INFγ^ or M^IL4^) (Supplementary Figure 2) were exposed to simvastatin (6 ng/ml) or atorvastatin (30 ng/ml, 70 ng/ml) for 24 hours. Macrophage scavenger receptor genes (MRC1, OLR1, MSR1, CD36 and SCARB1) were analysed by RT-qPCR pre- and post-exposure were measured.

We found that a 24-hour exposure to simvastatin or atorvastatin did not significantly alter the expression of these macrophage-specific genes.

## Discussion

### Different Macrophage Subsets in Human Carotid Atherosclerotic Plaques

Stroke is the second leading cause of death and one of the major causes of disability and dementia worldwide [33]. Around 80% of all stroke cases are ischaemic stroke, and of those, up to 20 % are due to large artery atherosclerosis (fatty blockage, often within the carotid artery) [34] [35]. Stroke survivors with a high atherosclerotic burden and unstable plaque morphology, e.g., inflammation or high cellular content, are at increased risk of early recurrent ischaemic events [2]. In this study, two patients had suffered from multiple previous stroke/TIAs.

Since macrophages play a crucial role in the pathogenesis and resolution of inflammation after stroke [3, 4], we sought to characterise macrophage phenotypes by simultaneous visualisation of both proinflammatory (M1-like) and anti-inflammatory (M2-like) macrophages *in situ*.

The macrophage population in carotid plaques is remarkably heterogeneous and influenced by the microenvironment [36]. Exposure to certain medications, such as lipid-lowering or anti-inflammatory drugs, may affect their functional phenotype [4]. If it were possible to modulate disease-enhancing phenotypes, this may provide a therapeutic avenue to lower the risk of further ischaemic events. Therefore, after plaque classification and macrophage characterisation, we determined the effect of statins and glucocorticoids *in vitro* on human macrophages isolated from blood (from healthy adult volunteers and stroke patients) and macrophages derived from human carotid atherosclerotic plaques ex vivo.

Studies have described the cellular and molecular interactions in atherogenesis; however, our understanding of the processes involved is continuously developing [37]. One of the main cellular components of atherosclerotic plaques are macrophages, which are part of the inflammatory response. In this study, macrophages were abundant in the collected plaques, which is in keeping with the fact that macrophages participate in all stages of plaque formation and progression [3].

The proinflammatory M1-like (CD86 single-positive: CD68+CD86+MRC1-) macrophages are atherogenic and have a role in endothelial activation and oxidative stress. These macrophages enhance plaque stability by weakening the fibrous cap and enlarging the necrotic core [38]. We found that the M1-like pro-inflammatory macrophage prevalence (17%) was significantly higher in unstable plaque sections, while all the other phenotypes were more prevalent in stable plaque sections. These findings agree with the fact that M1 macrophages enhance plaque instability as they secrete proinflammatory cytokines, such as IL-1β, IL-7, and IL-8 [39]. The representation of M1-like macrophages was also significantly higher in the AHA type VI- (complicated) plaques than in the AHA type V- (late) plaques. M1-like macrophages were also proportionately more prevalent in plaque types characterised by a bigger lipid core, calcification, rupture (type VIa) and haemorrhage (type VIb), in keeping with previous studies [36, 39, 40].

The anti-inflammatory M2-like macrophages (MRC1 single-positive: CD68+CD86-MRC1+) are atheroprotective, have a regulatory function and can carry out collagen synthesis [41]. These macrophages promote neoangiogenesis and produce TGF-β and an immunosuppressive cytokine, IL-10 [38]. We found that M2-like anti-inflammatory macrophages were less abundant overall, accounting for 9% of all CD68+ macrophages. Whilst this proportion was observed through all AHA-classified plaque sections, the anti-inflammatory (M2-like) macrophage phenotype was significantly lower than the pro-inflammatory (M1-like) macrophages in the plaque shoulder regions.

Several other studies support these data [38]. It was previously reported that the relative proportion of M2 macrophage phenotype is low in symptomatic plaques [36, 42] and that these are mainly found in the adventitia layer [10]. It was also reported that M2 macrophages dominate early atherosclerotic plaques, whereas advanced plaques contain mostly M1 subtypes [38], which is supported by our findings. The fact that the M1 and M2 macrophage ratio is different in early and advanced plaques could support the concept that macrophages can become altered by their microenvironment and change phenotype based on the surrounding signals.

The double-positive macrophages (hybrid M1 and M2) accounted for the majority (68%) of all CD68+ cells throughout all AHA-classified plaque sections. This macrophage subtype also represented 27% of all carotid artery plaque macrophages in the cite-seq dataset of Fernandez [28]. Furthermore, these cells were clustered, meaning these cells are more similar to each other than those in different clusters. These macrophages may, therefore, represent a previously undescribed macrophage subpopulation that warrants further exploration in future studies.

The double-negative (neither M1 nor M2) macrophage population was the smallest subset (7%) in carotid plaques. Interestingly, the proportion of this previously uncharacterised macrophage phenotype was higher in the stable plaque sections. When we re-analysed the dataset by Fernandez [28], we identified that 13% of macrophages were “double-negative” and were scattered across all macrophage groups. Therefore, these could be ‘intermediates’, perhaps representing macrophages that are in the process of switching from one phenotype to another, supporting that macrophage functions can change in disease or response to external stimuli [43].

### The effect of glucocorticoids on macrophage-specific genes in human polarised macrophages

In medicine, glucocorticoids are widely used as anti-inflammatory and immunosuppressive medications. Glucocorticoid hormones regulate fundamental physiological functions, like stress response and energy homeostasis [16]. Anti-inflammatory synthetic glucocorticoids such as dexamethasone suppress nuclear factor kappa B (NFκB) and downregulate proinflammatory cytokines (IL-1, IL-6, TNFα) [13] [44]. For example, GCs can inhibit vascular permeability and endothelial activation by directly inhibiting endothelial adhesion and indirectly inhibiting cytokine transcription [45].

In terms of specific agents, dexamethasone (0.25 mg/kg) reduces infarct volume size, mortality and neurological deficits in a mouse ischaemic stroke model [20]. Moreover, dexamethasone-treated (100 nM) macrophages were found to be more resistant to LPS-induced apoptosis [44]. Dexamethasone (100 nM for 48h) also promotes macrophage survival to decrease inflammation through its anti-apoptotic effects [46]. In addition, lower doses of GCs (corticosterone) increased NO production, while higher doses of GCs reduced NO production in macrophages [47], suggesting a dose-dependent effect.

In this study, we report that the expression of macrophage-specific genes, including scavenger receptors in M^LPS+INFγ^ and M^IL4^ macrophages, was altered following 24-hour exposure to different, clinically relevant concentrations of dexamethasone. No such alteration occurred following exposure to statins, despite statins being thought to reduce cardiovascular event risk via anti-inflammatory effects as well as lipid-lowering effects[48]. Similarly, in another study of the monocytic THP-1 cell line, there was no effect of atorvastatin treatment on the expression of macrophage scavenger receptors SR-AI (*MSR1*) or *CD36* [49].

Others have demonstrated that dexamethasone (0.3 µM= 117.65 ng/ml) upregulated *MRC1* gene expression (up to 100 fold) in human polarised pro- and anti-inflammatory macrophage cell line (THP-1) [50]. We have further shown that at clinically relevant concentrations, as low as 2 ng/ml, dexamethasone re-programs macrophages isolated from the blood of healthy volunteers and stroke patients to a less inflammatory state by increasing *MRC1* gene expression.

These results indicate that dexamethasone may have an anti-inflammatory effect on human macrophages through the induction of MRC1, an inflammatory mediator [51].

In our study, the average *OLR1* gene and protein expression was significantly reduced following exposure to dexamethasone in hMDMs isolated from healthy volunteers and stroke patients. These results align with Hodrea’s [52] earlier findings that *OLR1* gene expression was reduced in monocyte-derived human dendritic cells after dexamethasone (10 nM-100 nM) treatment [52]. Cortisol, another glucocorticoid (0.1 to 1000 nM), has also been shown to reduce oxLDL uptake in macrophages differentiated from THP1 cells [53].

OxLDL accumulation by dexamethasone-treated hMDMs was reduced by around 50% in our functional tests, implying that M^LPS+INFγ^ and M^IL4^ macrophages bound less oxLDL. If such an effect were to occur *in vivo*, in patients taking dexamethasone, this could potentially lead to reduced formation of atherosclerotic plaque in vessel walls. In support of this concept, previous studies have shown that dexamethasone significantly decreased the development of aortic atherosclerotic plaque in rabbits by reducing the number of macrophages and T lymphocytes and inhibiting LDL uptake [17].

Furthermore, the effect of dexamethasone on human atherosclerotic plaques *ex vivo* was also examined. IF stain of the treated dexamethasone-treated carotid plaques showed that MRC1 expression of CD68+ plaque macrophages was increased by 87%, and the OLR1 expression was decreased by 54%. A single-cell suspension was made from the cultured carotid plaques to study the result of the dexamethasone treatment on the plaque as a whole. The effect seen was similar to those observed in our *in vitro* macrophage experiments in that MRC1 expression was increased 2.5-fold, and OLR1 expression was reduced 0.8-fold.

While these observations highlight the acute effects of GCs on plaque macrophage morphology and possible short-term protective effects, the long-term effects on plaques remain unclear. It is known that extended GC administration can elevate cardiovascular risk, potentially by accelerating atherothrombotic progression [54]. Therefore, investigating the prolonged effects of GCs on plaques is also necessary.

### The lack of effect of statins on macrophages specific genes in human M1 and M2a polarised cells

Our finding that exposure to statins did not alter macrophage phenotype as measured by MRC1 and OLR1 gene expression suggests that if statins do exert their therapeutic effects in cardiovascular risk reduction via anti-inflammatory mechanisms, these are by different mechanisms to those we observed with dexamethasone. Since statins are currently part of the best medical treatment for the majority of patients with recently unstable atherosclerotic plaque events, the addition of an anti-inflammatory agent, such as dexamethasone, with a different mechanism of action, could, therefore, provide additional benefits in terms of event risk reduction. However, this would need to be tested further in clinical studies.

### Conclusion

To conclude, we determined that M1-like macrophages are more prevalent in unstable carotid plaque regions. There was also a high prevalence of macrophages in transitional states. Polarised human macrophages displayed increased *MRC1* expression and decreased *OLR1* expression following exposure to clinically relevant concentrations of glucocorticoids. This was demonstrated in blood samples from both healthy human volunteers and from stroke survivors moreover in carotid plaque tissue cultured *ex vivo* from patients who underwent endarterectomy for recently symptomatic carotid stenosis. On further testing, dexamethasone exposure led to reduced Oxidised LDL uptake in carotid plaque cells cultured *ex vivo*. These results suggest that human macrophages can be reprogrammed to a less pathogenic state *in vitro* and *ex vivo* and provide a potential mechanism by which dexamethasone may help to stabilise atherosclerotic plaques in humans [44].

In the future, the *in vitro* polarised macrophage model we have developed here could be used as a mechanistic marker of glucocorticoid function in atherosclerosis. Moreover, clinical trials in patients with advanced atherosclerosis could evaluate anti-inflammatory agents (such as dexamethasone) to stabilise plaques and reduce the risk of recurrent ischaemic events.

However, there were some potential limitations. Only a limited number of genes and proteins involved in atherosclerosis were examined. Additionally, macrophage activation states are better represented by a spectrum model rather than the binary M1 versus M2 polarization model [55]. The sample sizes were also small, although the effect of dexamethasone treatment was consistent across all the human tissue and blood samples.

## Nonstandard Abbreviations and Acronyms

GC: glucocorticoid
h.cort: hydrocortisone
H&E stain: haematoxylin and eosin stain
hMDMs: human monocyte-derived macrophage
EVG stain: Elastin van Gieson
IL4: Interleukin 4
INF-γ: interferon-gamma
LDL: low-density lipoprotein
LPS: lipopolysaccharide
M-CSF: macrophage colony-stimulating factor
oxLDL: oxidised low-density lipoprotein
PBMC: peripheral blood mononuclear cell
pred: prednisolone
VLDL: very low-density lipoprotein

## Acknowledgements

We gratefully acknowledge the assistance of the departmental technical team, the microscopy lead, the vascular surgery team and all the nurses who helped us with our work. A special thanks go to all volunteers and patients who donated their blood or tissue for research.

