## Supplementary for "Reprogramming Human Inflammatory Macrophages in Symptomatic Carotid Stenosis: Potential Mechanisms for Stabilisation of Atherosclerotic Carotid Plaques"

### Methods

#### *Governance*

All experiments were performed in accordance with United Kingdom legislation under the Human Tissue Act 2004. Carotid plaques were collected from The Northern General Teaching Hospitals in Sheffield in accordance with the study protocol (REC reference 14/SC/0147). Peripheral blood mononuclear cells were isolated from healthy adult volunteers in accordance with ethics approved by the University of Sheffield ethics committee (031330). According to Good Clinical Practice Guidelines, all patients and volunteer blood donors were given the information sheet and time to read through and ask any questions before informed consent was obtained.

#### *Histological processing and assessment of human carotid plaques*

Carotid plaques (n=13for the histology study) were removed from patients undergoing carotid endarterectomy and placed in 10% (v/v) neutral buffered. The carotid plaque was decalcified in an excess 0.5 M EDTA at pH 8 for 7 days and sectioned as described by Lovett et al [7, 8]. Overall stability [7, 8] and the American Heart Association grade [26,]was measured based on different histological features such as haemorrhage, lipid core, calcification, rupture, plaque macrophages, and cap inflammation. Histology was performed by KS or JR after H&E, EVG (ab150667), CD3 (A0452; MP-7401) and CD68 staining (M0876; MP-7401).

#### *Immunofluorescence staining of atherosclerotic plaques*

FFPE atherosclerotic plaque sections were de-waxed in xylene (Fisher Scientific) and rehydrated to water through graded ethanol (Fisher Scientific). Endogenous peroxide activity was blocked with 3% (v/v) $H_2O_2$ (Merck) in methanol (Fisher Scientific) for 10 minutes. Heat-mediated antigen retrieval was performed in antigen unmasking solution (H-3300, Vector) to break down cross-links between antigens. Non-specific binding was blocked by incubating the sections in animal serum-free protein blocking reagent (GTX30963, GeneTex) for 60 minutes at room temperature. Sections were incubated with primary and then with secondary antibodies, summarised in **(Supplementary Table 1)**.

ProLong Gold Antifade Mountant with 4′,6-diamidino-2-phenylindole (DAPI) (Thermo Fisher Scientific) was used to mount the sections and counterstain the nuclei. Images were captured using a Leica AF6000LX inverted microscope.

Negative and positive controls were added to ensure specific stains. Purified mouse (Abcam) or rabbit (Vector Laboratories) IgG were used in the same concentration as primary antibodies for negative control. Human tonsil tissue was used as a positive control for CD68 antibody, while the human spleen and lung were used for CD86 and MRC1 antibodies, respectively, as suggested by manufacturers.

*Supplementary Table 1: Summary of primary and secondary antibodies*

<table>
<tr><th colspan="2"></th><th>Primary antibody</th><th>Secondary antibody</th></tr>
<tr><td rowspan="3">Different macrophage subsets in carotid plaques</td><td>Single IF staining</td><td>Mouse anti-human CD68 (ab125157) 1:50</td><td rowspan="2">Donkey anti-mouse NL493 (NL009) 1:400</td></tr>
<tr><td rowspan="2">Dual IF staining</td><td>Mouse anti-human CD86 (ab213044) 1:50</td></tr>
<tr><td>Rabbit anti-human MRC1 (ab64693) 1:50</td><td>Donkey anti-mouse NL557 (NL004) 1:400</td></tr>
<tr><td rowspan="4">Ex vivo carotid plaque culture and drug treatment</td><td rowspan="2">Dual IF staining</td><td>Mouse anti-human CD86 (ab213044) 1:50</td><td>Alexa 488 anti-mouse (4408) 1:400</td></tr>
<tr><td>Rabbit anti-human OLR1 (ab126538) 1:50</td><td>Alexa 594 anti-rabbit (8889) 1:400</td></tr>
<tr><td rowspan="2">Dual IF staining</td><td>Mouse anti-human CD86 (ab213044) 1:50</td><td>Alexa 488 anti-mouse (4408) 1:400</td></tr>
<tr><td>Rabbit anti-human MRC1 (ab64693) 1:50</td><td>Alexa 594 anti-rabbit (8889) 1:400</td></tr>
</table>

***Image Analysis***

For the macrophage subsets experiments, regions of interest (plaque shoulder regions) were analysed across sections with FIJI. The percentages of macrophages (CD68+) which were single positive (CD86+MRC1- or CD86-MRC1+), double positive (CD86+MRC1+) or double negative (CD86-MRC1-) were calculated.

During the ex vivo carotid plaque analysis studies, the IF intensity and percentage of OLR1 or MRC1 positive macrophages were calculated with FIJI.

***Peripheral Blood Mononuclear Cell (PBMC) isolation***

Whole blood was taken by venepuncture from human subjects and anticoagulated with 3.8 % (w/v) trisodium citrate dihydrate (Fisher Scientific). After mixing by inversion, whole citrated blood was layered carefully onto Ficoll-Paque Plus (GE Healthcare, Life Sciences) in a 2:1 ratio. After overlaying the blood, the tube was centrifuged at 900 g, at room temperature, for 20 minutes with acceleration and brake set at 1. The layer of PBMCs was collected in 5 ml phosphate-buffered saline- EDTA (2mM) (PBSE). Cells were then centrifuged at 400 g, at room temperature, for 5 minutes. The resulting soft cell pellet was resuspended in 10 ml red blood cell (RBC) lysis buffer (155 mM $NH_4Cl$, 10 mM $KHCO_3$, 0.1 mM EDTA) and incubated for 5 min at room temperature. The suspension was topped up to 50 ml with PBSE and centrifuged at 400 g at room temperature for 5 minutes.

***CD14+ monocyte isolation***

The PBMC suspension was centrifuged at 400 g at room temperature for 5 minutes. The PBMC pellet was resuspended in 90 µl cold MACS buffer (0.5% (w/v) bovine serum albumin (BSA) in PBSE), followed by the addition of 10 µl CD14 microbeads (Miltenyi Biotec) per 10 million PBMCs and incubation for 15 minutes at 4 °C. A further 2 ml MACS buffer was added, and the solution was centrifuged at 300 g at room temperature for 5 minutes. The cell pellet was resuspended in 500 µl MACS buffer. Magnetic separation was carried out by LS columns (Miltenyi Biotec), and CD14+ cells were collected.

***Macrophage differentiation***

The CD14+ cell suspension was centrifuged at 400 g at room temperature for 5 minutes. The cell pellet was resuspended in complete media at the desired cell density and cultured at 37°C with 5% $CO_2$. The culturing media (RPMI-1640, Gibco) was supplemented with 10% (v/v) ultra-low endotoxin heat-inactivated fetal bovine serum (FBS) (Pan Biotech), 1% (v/v) L-glutamine (Gibco), 1% (v/v) Penicillin-Streptomycin (Gibco). MDMs were differentiated in this complete media on plastic wells with the addition of 100 ng/ml recombinant human Macrophage Colony Stimulating Factor (rh M-CSF, Gibco) for 7 days. PBMC, CD14+ monocyte isolation and macrophage differentiation protocols were developed based on a previously established method by Geng and Hansson.

***Macrophage polarisation***

After differentiation (Day 7), the polarisation of $M^{LPS+INF\gamma}$ and $M^{IL4}$ macrophages was performed by culturing cells for 24 hours in fresh, complete media supplemented with specific polarising factors. Cells destined towards $M^{LPS+INF\gamma}$ macrophages were polarised by LPS (100 ng/ml, Enzo Life Sciences) and interferon-gamma (INF-γ) (20 ng/ml, Pepro Tech). Cells destined towards the $M^{IL4}$ phenotype were polarised using IL-4 (20ng/ml, Pepro Tech). In each of the following assays, unpolarised macrophages ($M^{un}$) were used as controls.

***Justification for concentrations of dexamethasone used***

During previous pharmacokinetic studies, 10 healthy female volunteers received 0.5 mg oral administration (p.o.) or 1.5 mg p.o. dexamethasone. After taking the appropriate dose of dexamethasone, blood samples were taken at 10 different time points. The elimination of dexamethasone was rapid, and after 24h, the plasma concentration was less than 1/10$^{th}$ of the maximal concentration in all dose groups. At 0.5 mg p.o. dexamethasone dose, the average plasma level was 8 ng/ml, while after 1.5 mg p.o. dexamethasone, the plasma level reached 14 ng/ml. Similar results were reported where 98.1 ng/ml serum level was measured after 7.5 mg p.o. dexamethasone (n=54). We therefore selected 8, 14, and 125 ng /ml. When it became apparent that all of these doses had an effect on macrophage phenotype, we tested two further (lower) concentrations of 2 and 4 ng/ml.

***Supplementary Table 2: Dose-dependent pharmacokinetics of dexamethasone measured by radioimmunoassay***

*Plasma level ($C_{max}$) is shown as an average amongst n=10 healthy adults after a single dose (p.o.) of dexamethasone.*

| **Name** | **Dosage** | **Plasma level ($C_{max}$) (Average of n=10)** |
|---|---|---|
| **dexamethasone I** | 0.5 mg | 8 ng/ml |
| **dexamethasone II** | 1.5 mg | 14 ng/ml |
| **dexamethasone III** | 10 mg | 125 ng/ml |

***Macrophage treatment with statins and glucocorticoids***

Macrophage treatment with clinically relevant statin concentrations was performed on day 7 by culturing cells for 4 hours in fresh complete media supplemented with atorvastatin (30 ng/ml, 70 ng/ml, Merck) or simvastatin (6 ng/ml, Merck). After this, cells were polarised towards either $M^{LPS+INF\gamma}$ or $M^{IL4}$ phenotype in addition to statins for 24 h.

Macrophage treatment with dexamethasone (Merck) was performed in fresh complete media supplemented with different clinically relevant concentrations of dexamethasone (8 ng/ml, 14 ng/ml or125 ng/ml), or the two lower concentrations of dexamethasone (2 ng/ml or 4 ng/ml) or with 5.2 µg/ml prednisolone (Merck) or 2.8 µg/ml hydrocortisone (Merck). After 24h incubation, cells were washed twice with PBS 1X and prepared for further experiments.

***RNA isolation***

RNA was isolated using RNeasy UCP Micro Kit (Qiagen) according to the manufacturer's instructions. RNA concentration was measured using spectrophotometric analysis (Nanodrop), measuring the absorbance of each sample at 260nm.

***cDNA synthesis and Real-Time quantitative PCR***

cDNA synthesis was performed using the iScript cDNA synthesis kit (Biorad) according to the manufacturer's instructions. This kit operates up to 15 µl of RNA sample in a 20 µl cDNA synthesis reaction.

RT-qPCR was performed using a Precision PLUS qPCR Master Mix with SYBR Green (Primer Design), and results were analysed with CFX384 C1000 Touch Thermal Cycler (Biorad). Primer sequences and qPCR cycle conditions (**Supplementary Table 3)**.

***Supplementary Table 3: Real-Time qPCR cycle conditions (top) and primers (bottom) for testing of human genes***

| Steps | Settings |
|---|---|
| **Polymerase activation Denaturation** | 95°C; 2 min |
| **Amplification** | 40 amplification cycles:<br>95°C; 15 secs<br>60°C; 1 min |
| **Melting Curve Analysis** | 60°C- 95°C/ 0,5°C Increment |

| Name | Sequence | Length (nt) | Tm (°C) | GC % |
|---|---|---|---|---|
| **GAPDH Fwd** | ATTGCCCTCAACGACCACTTT | 21 | 52 | 48 |
| **GAPDH Rev** | CCCTGTTGCTGTAGCCAAATTC | 22 | 55 | 50 |
| **CD68 Fwd** | AAGGGGGCTCTTGGGAACTA | 20 | 54 | 55 |
| **CD68 Rev** | CCAAGCCCTCTTTAAGCCCC | 20 | 56 | 60 |
| **CD86 Fwd** | CCCAGACCACATTCCTTGGAT | 21 | 54 | 52 |
| **CD86 Rev** | TCCCTCTCCATTGTGTTGGT | 20 | 52 | 50 |
| **MRC1 Fwd** | CCATCGAGGAAGAGGTTCGG | 20 | 56 | 60 |
| **MRC1 Rev** | GGGTGGGTTACTCCTTCTGC | 20 | 56 | 60 |
| **OLR1 Fwd** | TGCGACTCTAGGGGTCCTTTG | 21 | 56 | 57 |
| **OLR1 Rev** | TGTTAGGAGGTCAGACACCTGG | 22 | 57 | 55 |
| **MSR1 Fwd** | CGAGGTCCCACTGGAGAAAGT | 21 | 56 | 57 |
| **MSR1 Rev** | CAATTGCTCCCCGATCACCTTT | 22 | 55 | 50 |
| **CD36 Fwd** | TCTGTCCTATTGGGAAAGTCACTG | 24 | 56 | 46 |
| **CD36 Rev** | GAACTGCAATACCTGGCTTTTCTC | 24 | 56 | 46 |
| **SCARB1 Fwd** | GAATCCCCATGAACTGCTCTGT | 22 | 55 | 50 |
| **SCARB1 Rev** | TCCCAGTTTGTCCAATGCCTG | 21 | 54 | 52 |

***Protein analysis with Western Blot***

Polarised cells were lysed overnight in radioimmunoprecipitation assay (RIPA) buffer (Merck) in the presence of phosphatase inhibitors (PhosSTOP, Roche) and protease inhibitors (Protease Inhibitor Cocktail, Merck) as per the manufacturer's instruction. A BCA Assay was used to quantify total protein before western blotting.

15 µg of sample protein was added to NuPage LDS Sample Buffer (Invitrogen) up to a final volume of 20 µl and incubated at 70°C for 10 minutes. Equal amounts of protein sample were loaded into the wells of a NuPAGE 4-12% Bis-Tris SDS-PAGE Gel (Invitrogen) together with a standard protein size marker (SeeBlue, Plus2 Prestained Standard, Invitrogen). Electrophoresis was performed at 120V for 70 minutes in NuPAGE MES SDS Running Buffer (Novex, Invitrogen). Proteins were then transferred onto a nitrocellulose membrane at 35 V for 60 minutes in NuPAGE Transfer Buffer (5% (v/v) 20x transfer buffer, 0.1% (v/v) antioxidant, 20% (v/v) methanol. NuPAGE Antioxidant (Novex, Invitrogen) was added to the Transfer Buffer (0.1% (v/v)) to enhance the transfer of proteins to membranes. The membrane was blocked at room temperature for 1 hour in 5% (w/v) milk TBST. The membrane was incubated at 4°C overnight with primary antibody (OLR1, 1:500, Abcam) diluted in 5% (w/v) milk TBST. After incubation, the membrane was washed 3 times for 5 minutes in TBST and incubated with the horseradish peroxidase (HRP) conjugated secondary antibody (1;2500, Agilent) diluted in 5% (w/v) milk TBST at room temperature for 1 hour. The membrane was washed 3 times for 5 minutes in TBST.

Specific protein bands were detected by adding a chemiluminescent substrate (SuperSignal™ West Pico PLUS Chemiluminescent Substrate, Thermo Fisher) and using the ChemiDoc XRS+ Imaging System (Biorad).

In order to use different primary antibodies (i.e., housekeeping: GAPDH, 1:5000, Santa Cruz Biotechnology), the membrane was stripped in ReBlot Plus Strong Antibody Stripping Solution 1X (Millipore) for 20 minutes at room temperature. The membrane was re-blocked at room temperature for 1 hour in 5% (w/v) milk TBST before using new primary and secondary antibodies.

Results were analysed in Excel through Image Studio Lite version 5.2.5. First, the intensity for each protein band was calculated with background subtraction. The intensity was normalised for each test sample (adjusted intensity $_{\text{protein of interest}}$ / adjusted intensity $_{\text{housekeeping}}$).

***OxLDL accumulation in MDMs, Oil Red O Staining, microscopy and analysis***

To detect the oxLDL accumulation in macrophages, human MDMs were plated and polarised in 12-well plates. After 1 day of polarisation in the presence of dexamethasone, cells were washed 3 times with PBS 1X and human oxLDL was added for 24h (working concentration: 25 µg/ml; Thermo Fisher).

Oil Red O (ORO) staining (according to the manufacturer's instructions, Merck) was used to detect lipid accumulation in hMDMs. After oxLDL accumulation, cells were fixed with 4% (w/v) PFA (Merck). Before staining, cells were rinsed with water and 60% (v/v) isopropanol. 1 ml of ORO working solution was added to the cells for 15 mins, then removed and rinsed with 60% (v/v) isopropanol and washed at least 3 times with distilled water. Haematoxylin ((Fisher Scientific) counterstain was used, and cells were observed with a LeicaDMI4000B inverted microscope with Leica Application Suite Advanced Fluorescence (LASAF) v2.63 software (Microscopy Core Facility) was used.

The percentage of positive cells was counted, and quantifications were performed on an average of 5 fields of view.

***OLR1 IF stain, microscopy and analysis***

Human MDMs were plated and polarised in 8-well chamber slides (Thermo Fisher) and stained to detect the OLR1 protein, which binds oxLDL. Immediately after polarization, cells were fixed with 4% (w/v) PFA at room temperature for 30 minutes, washed 3 times with PBS 1X and stored at 4°C until further work.

Cells were permeabilised with 0.1% (v/v) Triton PBS for 15 minutes, washed 3 times with PBS 1X, then blocked with 2% (w/v) BSA-PBS for 45 minutes at room temperature and washed 3 times with PBS 1X. Cells were incubated with rabbit polyclonal OLR1 primary antibody (Abcam) (1:200) for 1 hour at RT. After washing 3 times with PBS 1X, a positive stain was detected by donkey anti-rabbit IgG-NL557 conjugated antibody (R&D Systems). ProLong Gold Antifade Mountant with DAPI (Thermo Fisher Scientific) was used to mount the sections and counterstain the nuclei.

Images were captured using Leica AF6000LX inverted microscope and with Leica Application Suite Advanced Fluorescence (LASAF) v2.63 software. OLR1 positive cells were counted and compared to the maximum cell number based on the DAPI IF signal and quantifications performed on an average of 5 fields of view.

***Ex vivo carotid plaque culture, treatment & digestion***

Four carotid plaques removed at carotid endarterectomy (CEA) were immediately placed in fresh media, and transferred on ice. The plaques were divided into 2 equal parts in cross way.

One part was dissected into smaller pieces (2x2mm) and cultured in complete media (RPMI) with and without dexamethasone (8 ng/ml) for 3 days at 37 °C in humidified air, containing 5% $CO_2$. To create a single cell suspension for RNA work, the RPMI media was supplemented with collagenase XI from Clostridium histolyticum (1.25 mg/ml, Merck) and deoxyribonuclease I (0.2 mg/ml, Merck). The suspension was incubated in a horizontal incubator shaker at 250 rpm for 1 hour 30 min at 37ºC and filtered (40-µm filter). Cells were washed twice with PBS 1X and prepared for further experiments.

The other part was divided into 3 further sections for the following subgroups: Day 0, Day 3- untreated and Day 3- dexamethasone-treated. After the histological procession, the tissue was IF stained with CD68 pan macrophage marker, OLR1 and MRC1 antibodies. The staining protocol is detailed in the section entitled Immunofluorescence staining of atherosclerotic plaques

### Results

***Supplementary Table 4: Overall Plaque Instability grade (as defined by Lovett et al [7, 8]) and AHA classification of the 11 plaques that were sufficiently intact for histological examination.***

| Study ID | Section | Stability | AHA classification |
|---|---|---|---|
| 1 | A | Probably unstable | VIa: Rupture |
| | B | Definitely unstable | |
| | C | Definitely unstable | |
| | D | Probably unstable | |
| | E | Probably unstable | |
| 2 | A | Definitely unstable | VIb: Haemorrhage |
| | B | Definitely unstable | |
| | C | Definitely unstable | |
| | D | Probably stable | |
| 3 | A | Probably unstable | Va: Multilayer fibrous atheroma with core |
| | B | Probably stable | |
| | C | Probably stable | |
| | D | Probably unstable | |
| 4 | A | Probably unstable | VIa: Rupture |
| | B | Definitely unstable | |
| | C | Probably unstable | |
| | D | Probably unstable | |
| 5 | A | Probably unstable | VIa: Rupture |
| | B | Definitely unstable | |
| | C | Definitely unstable | |
| 6 | A | Probably stable | Va: Multilayer fibrous atheroma with core |
| | B | Definitely stable | |
| | C | Probably stable | |
| | D | Probably stable | |
| 7 | A | Probably stable | Va: Multilayer fibrous atheroma with core |
| | B | Probably stable | |
| | C | Probably stable | |
| | D | Probably stable | |
| 8 | A | Probably unstable | Va: Multilayer fibrous atheroma with core |
| | B | Probably unstable | |
| | C | Definitely unstable | |
| | D | Definitely unstable | |
| | E | Definitely stable | |
| 9 | A | Definitely stable | Vc: fibrotic without lipid core |
| | B | Definitely stable | |
| | C | Definitely stable | |
| | D | Definitely stable | |
| 10 | A | Definitely unstable | Vb: calcification |
| | B | Definitely unstable | |
| | C | Probably unstable | |
| | D | Probably stable | |
| | E | Probably stable | |
| 11 | A | Definitely stable | Vc: fibrotic without lipid core |
| | B | Definitely stable | |
| | C | Definitely stable | |
| | D | Definitely stable | |

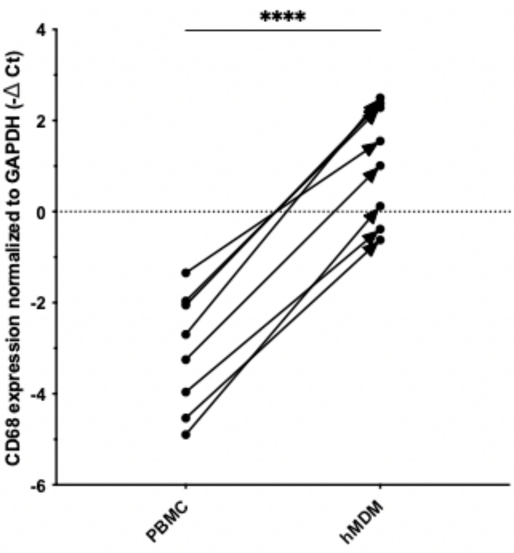


***Supplementary Figure 1: Validation of the differentiation of CD14+ PBMCs into hMDMs by the increased CD68 expression***

*The increased expression of CD68 and cell adhesion to the culture plate verified macrophage differentiation from PBMCs. The expression of the CD68 marker was significantly higher ($p < 0.05$) in hMDMs compared to the PBMCs. Data is plotted from 8 donors, individually listed (n=8), normalised to the GAPDH housekeeping gene. The direction of change in expression is shown as an arrow for each individual healthy volunteer. The dotted line indicates a 0-expression level. P-value: **** ‹0.0001.*

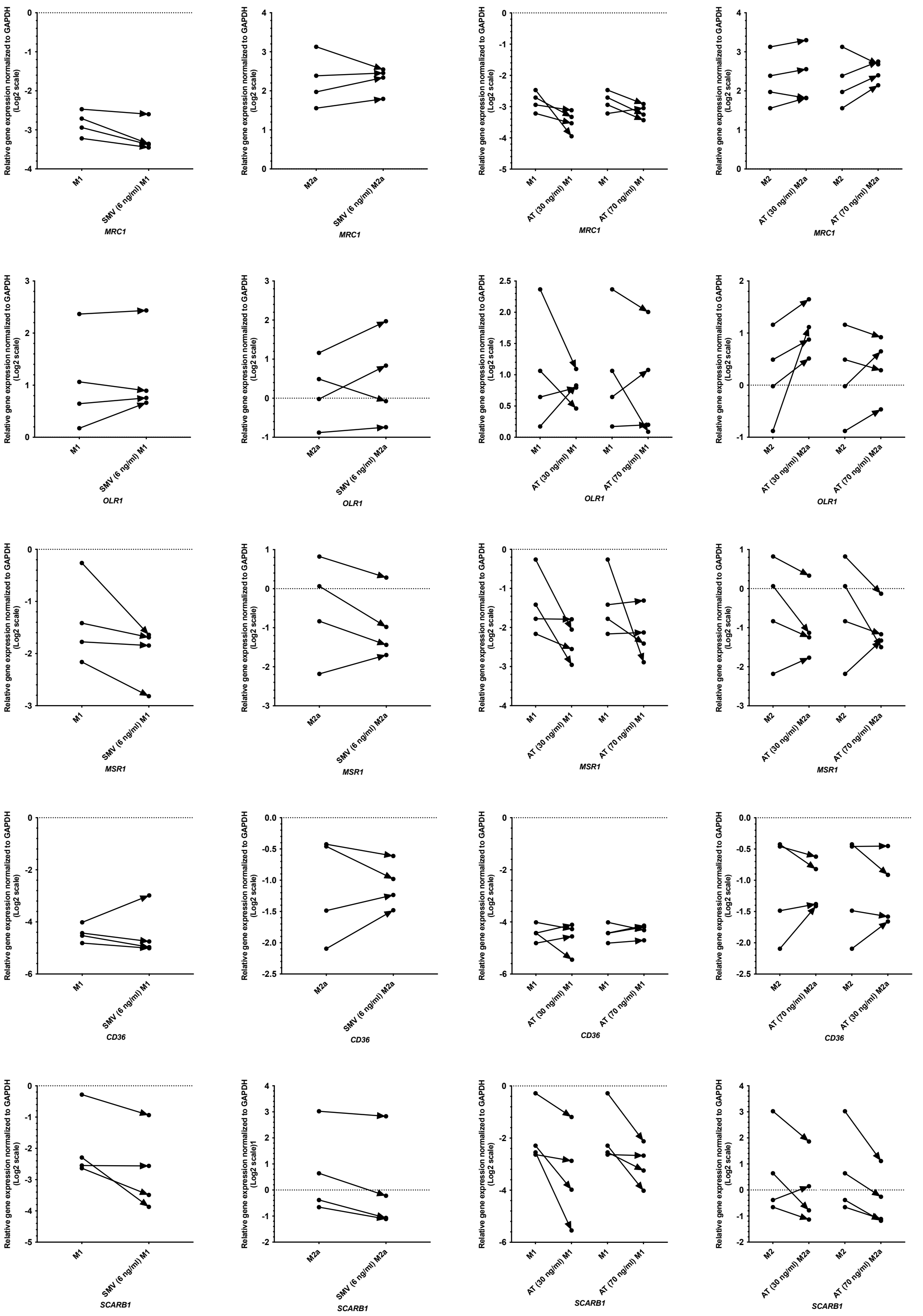


***Supplementary Figure 2: Relative gene expression analysis of MRC1, OLR1, MSR1, CD36, and SCARB1 in human MDMs after simvastatin and atorvastatin treatment.***

*Human* $M^{LPS+INF\gamma}$ *and* $M^{IL4}$ *polarised macrophages were treated with simvastatin (SMV) (6 ng/ml) or atorvastatin (AT) (30, 70 ng/ml)) and gene expression was determined with RT-qPCR. Data is plotted from healthy adult blood samples, individually listed (n=4), normalised to the GAPDH housekeeping gene. The direction of change in expression is shown with the treatment as an arrow for each individual donor. The dotted line indicates a 0 expression level.*

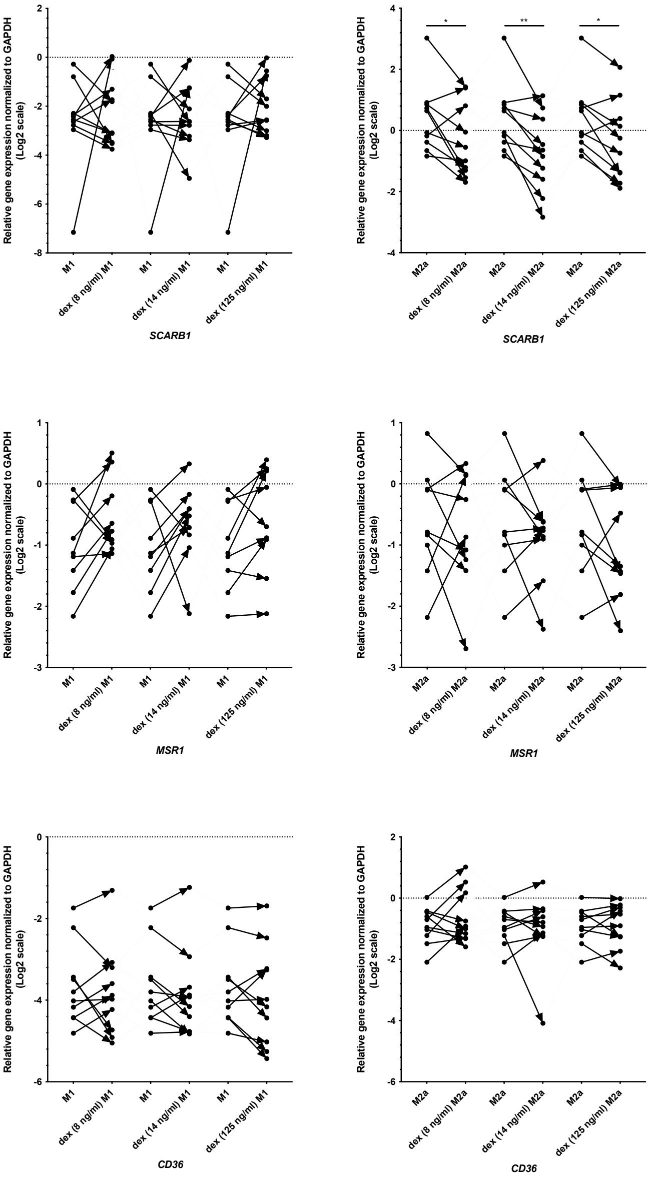


***Supplementary Figure 3: Relative gene expression analysis of SCARB1, MSR1 and CD36 in human MDMs after dexamethasone treatment***

*Human* $M^{LPS+INF\gamma}$ *(left panel) and* $M^{IL4}$ *(right panel) polarised macrophages were treated with dexamethasone (8, 14, 125 ng/ml), and SCARB1, MSR1 and CD36 gene expression was determined with RT-qPCR. Data is plotted from 9 healthy volunteer blood samples, individually listed and normalised to the GAPDH housekeeping gene. Data follow normal (Gaussian) distribution and are analysed with a two-tailed paired t-test. The direction of change in expression is shown with the treatment as an arrow for each individual donor. The dotted line indicates a 0 expression level. p-value: * ‹0.0332, ** ‹0.0021,*
